# UNCROSS2: identification of cross-talk in 16S rRNA OTU tables

**DOI:** 10.1101/400762

**Authors:** Robert C. Edgar

## Abstract

Next-generation amplicon sequencing is widely used for surveying biological diversity in applications such as microbial metagenomics, immune system repertoire analysis and targeted tumor sequencing of cancer-associated genes. In such studies, assignment of reads to incorrect samples (cross-talk) is a well-documented problem that is rarely considered in practice. Here, I describe UNCROSS2, an algorithm designed to detect and filter cross-talk in OTU tables generated by next-generation sequencing of the 16S ribosomal RNA gene. On eight published datasets, cross-talk rates are estimated to range from 0.4% to 1.5% mis-assigned reads. On a mock community test, UNCROSS2 identifies spurious counts due to cross-talk with sensitivity ∼80% to 90% and error rate from ∼1% to ∼20%, but it is not clear whether the accuracy of the algorithm is sufficient to decisively improve diversity rates in practice.

## Introduction

Recent examples of next-generation amplicon sequencing experiments include the Human Microbiome Project (HMP Consortium *et al.*, 2012), which sequenced the 16S ribosomal RNA (rRNA) gene, an analysis of the response of the human immune system to influenza vaccination (Jiang *et al.*, 2013), which sequenced antibody immunoglobulin genes, and a high-throughput search for cancer-relevant variants in 16 oncogenes (Hadd *et al.*, 2013). In such studies, samples are usually multiplexed by embedding index sequences into PCR primers which identify the sample of origin. Index sequences are sometimes called tags or barcodes, but I will avoid the latter terms here as some authors use them to refer to the biological sequence in an amplicon. An index sequence can be embedded in the forward primer (Caporaso *et al.*, 2011; Derakhshani *et al.*, 2016) (*single-indexing*), while *dual-index* schemes embed indexes in both primers (Kozich *et al.*, 2013; Derakhshani *et al.*, 2016) to enable larger numbers of samples. Reads are assigned to samples (*demultiplexed*) by identifying their index sequences. A *cross-talk* error occurs when a read is assigned to an incorrect sample. Previous studies have revealed unexpectedly high rates of cross-talk in both 454 (Carlsen *et al.*, 2012) and Illumina (Kircher *et al.*, 2012; Nelson *et al.*, 2014) data, but the causes of cross-talk are currently not well understood. Indexing methods designed to mitigate cross-talk have recently been proposed by (Esling *et al.*, 2015) and (Schnell *et al.*, 2015), but so far have rarely been used in practice. Here, I describe UNCROSS2, an algorithm designed to detect and filter cross-talk in 16S rRNA OTU tables.

## Methods

### Datasets

I analyzed eleven published datasets of 16S rRNA reads as summarized in Table 1. These studies sampled communities with low diversity (e.g. human vagina and prostate), moderate diversity (e.g. human gut) through high diversity (soil). In all but one of these datasets, as is typically the case in practice, samples were obtained from similar environments and there are no control samples with known composition. As an exception, the Koz2013 dataset includes samples from different environments (human gut, mouse gut and soil) together with designed (*mock*) control samples of known composition. Koz2013 contains reads from eleven MiSeq runs which were processed using a total of three different versions of the Illumina Real-Time Analysis (RTA) and MiSeq Control Software (MCS). Twelve samples were sequenced in each run: three replicates of a mock sample and three replicates obtained from human gut, mouse gut and soil, respectively.

**Table 1.**
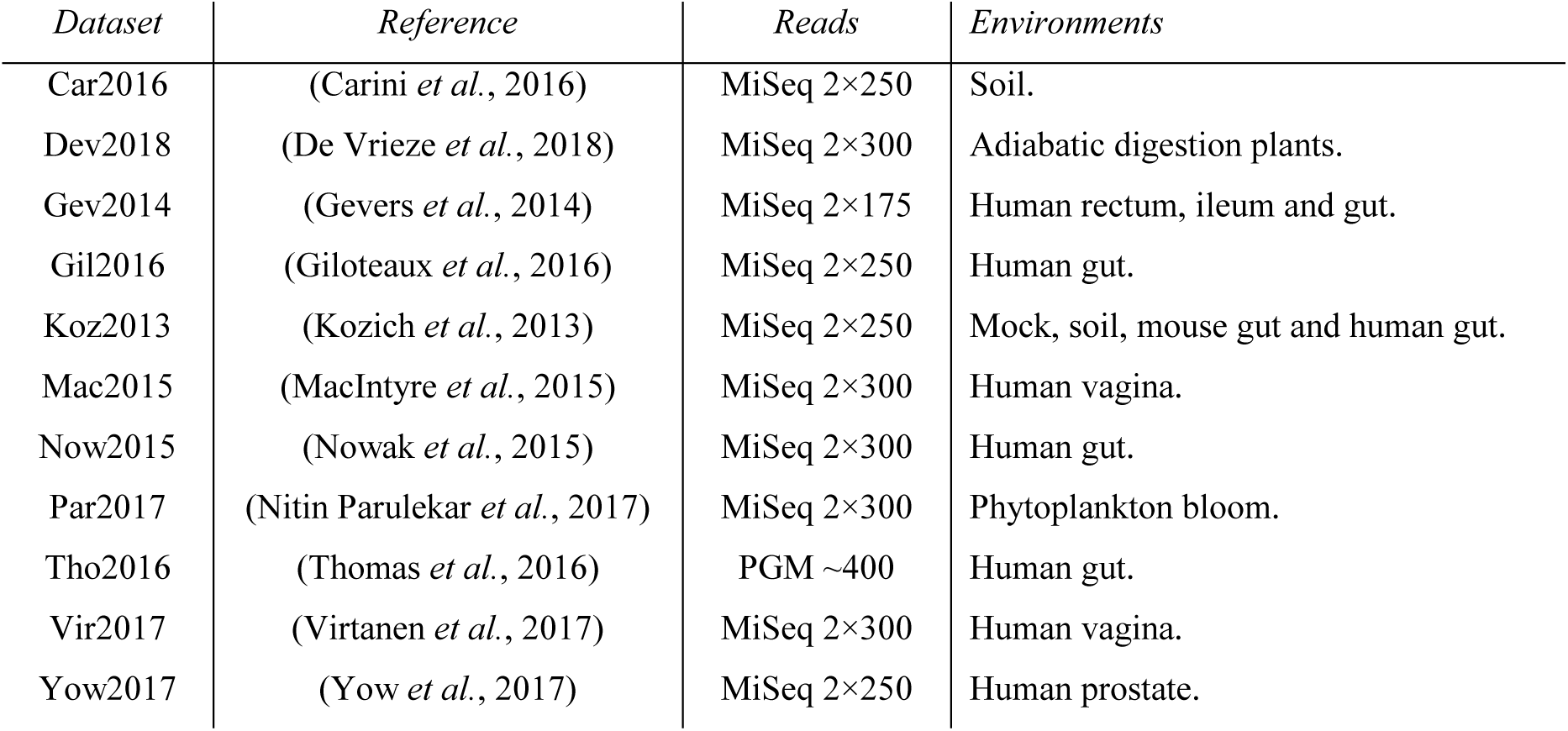
16S rRNA datasets analyzed in this paper.

### OTU tables

For all datasets, I generated OTU tables using the current recommended UPARSE (Edgar, 2013) protocol (https://drive5.com/usearch/manual/uparse_pipeline.html, accessed 1st August 2018).

### Cross-talk rate estimate with mock samples

The rate of cross-talk was estimated using mock samples as follows. Find all OTUs which do not match known sequences in the mock species and have non-zero read counts in both mock samples and other samples. The non-zero counts for mock samples in these OTUs are most likely explained by cross-talk (contaminants are also possible). Let *N* be the total number of reads in these OTUs, *M* be the total number of reads assigned to mock samples, *m* be the number of mock samples, and *n* be the total number of samples. Assuming that the total number of reads mis-assigned to each sample is approximately equal, the number of reads mis-assigned to all samples is *Mn*/*m*, and the fraction of mis-assigned reads overall is estimated to be

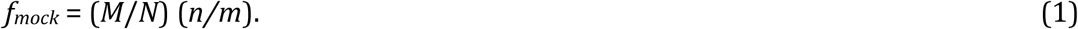

### De novo estimate of the cross-talk rate

When control samples are not available, the cross-talk rate must be estimated *de novo*, i.e. without prior knowledge of the sample composition. The UNCROSS2 algorithm estimates the rate by searching the OTU table for a subset of OTUs (*candidates*) which have the strongest evidence of cross-talk (Fig. 1). A candidate OTU has counts for some samples which are much smaller than the expected number that would be observed if the reads were evenly distributed over all samples. Such small counts are consistent with cross-talk, though this cannot be confirmed unless the composition of the sample can be determined independently of the OTU table. A count *c* is considered low if it is 0 < *c* ≤ *sN*_*i*_/*n*, where *N*_*i*_ is the total number of reads in the *i*th OTU and the user-settable parameter *s* is 0.1 by default so that a low count is at least ten times less than the mean for the OTU. Let *S*_*i*_ be the number of samples with low counts for this OTU, and *X*_*i*_ the sum of the low counts. The OTU is a candidate iff *N*_*i*_≥*N*_*min*_ and *S*_*i*_≥*S*_*min*_, where by default *N*_*min*_=1,000 and *S*_*min*_=3. A large value of *N*_*min*_ was chosen to avoid large random fluctuations which are likely to occur when counts are small, and *S*_*min*_ is greater than one as a check that the effect is reproduced in multiple samples. Low counts are tentatively inferred to be entirely due to cross-talk, and the estimated cross-talk rate *f*_*i*_ for a candidate OTU is calculated similarly to eq. (1) above,

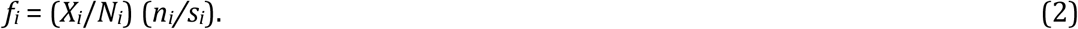

**Figure 1.**
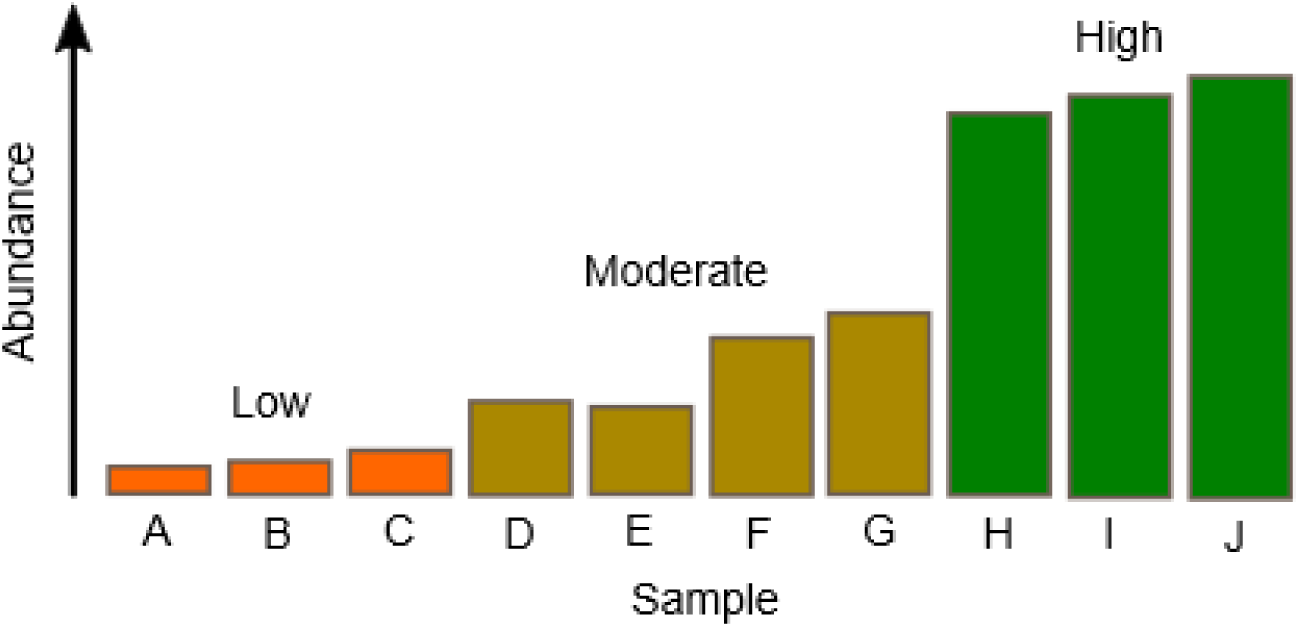
Abundance distribution of a candidate OTU. The histogram shows sample counts sorted in order of size. “Low” counts are much less than the mean, “moderate” counts are comparable to the mean, and “high” counts are much greater than the mean. A candidate OTU has at least three low counts which are greater than zero and less than 10% of the mean value. The low counts are tentatively interpreted as cross-talk.

If *f*_*i*_ is found to be greater than a plausible maximum for the cross-talk rate (*f*_*max*_, set to 0.02 by default), the OTU is rejected as a candidate. At least *C*_*min*_ candidate OTUs are required, with *C*_*min*_=10 by default; otherwise the data is considered insufficient to make a *de novo* estimate. To mitigate possible problems caused by false positives (FPs) and outlier cases, which could have anomalously high or low rates, the median rate for candidate OTUs is reported rather than the mean. The median rate is denoted *f* without a subscript.

### UNCROSS2 score

If the cross-talk rate is *f*, and the total number of reads assigned to the *i*th OTU is *N*_*i*_, then on average a count which should be zero will be *z*_*i*_=*f N*_*i*_/*n*, where *n* is the number of samples. Based on this observation, an algorithm could consider all counts ≤*z*_*i*_ to be consistent with cross-talk and set them to zero. However, this algorithm is likely to have a very high false negative (FN) rate, i.e. will fail to correctly identify many counts which are entirely due to cross-talk. First, the cross-talk rate between a given pair of samples may be much higher than the average rate. This will surely be the case for cross-talk due to base call errors in index reads. For example, if the correct indexes for a sample pair have a single difference, the rate will be much higher than a pair where all bases are different. Second, the rate will vary due to fluctuations, especially when counts are small. For example, suppose *f*=0.01, *n*=10 and *N*_i_=10. Then the expected value for a count which should be zero is *z*_*i*_=0.01, which is much less than one. But if we have 1,000 OTUs with *N*_*i*_=10, and the true count for all these OTUs is zero in in sample *A*, then we expect ∼10 of them to have a spurious non-zero count for *A*. In these OTUs, the observed cross-talk rate is at least 0.1 (one of ten reads is mis-assigned), which is much greater than the mean rate *f*. For OTUs where the cross-talk rate is much greater than *f*, a threshold of *z*_*i*_ is much too low. With these considerations in mind, I designed an *ad hoc* score *t* for a count *c* ranging from zero (minimum indication of cross talk) to one (maximum indication of cross-talk),

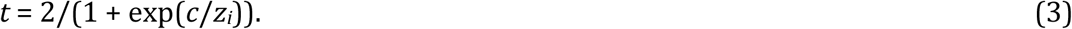

Here, exp(*x*) is the exponential function *e*^*x*^. If *c* is much less than the expected value *z*_*i*_, then *t* is close to 1, and if *c* is much greater than *z*_*i*_ then *t* is close to zero. The filtering threshold is specified as a minimum value of *t*, which by default is set to *t*_*min*_=0.1. This value is intended to identify a large majority of spurious counts due to cross-talk, at the possible expense of having a high FP rate, i.e. setting many counts to zero that should be at least one. This is because the most likely motivation for using a cross-talk filter is to improve estimates of alpha and beta diversity. A threshold which is designed to minimize the number of errors by balancing FPs and FNs, or to minimize FPs, is likely to leave many unfiltered spurious counts in the table. This is a similar situation to denoising algorithms, which set thresholds designed to minimize FNs (bad sequences which are falsely reported as correct) at the possible expense of a high rate of FPs (correct sequences which are falsely reported as bad) (Edgar, 2017). The value of denoising is undermined if the minority of bad sequences that remain after filtering are comparable to or more numerous than the correct sequences, which could easily happen given that the diversity of bad sequences is likely to be much larger than the diversity of correct sequences even if the base call error rate is very low (https://drive5.com/usearch/manual/tolstoy.html). With cross-talk, there is similarly little point in removing many or most spurious counts if the minority that remain after filtering could be more numerous than the valid non-zero counts. Both denoisers and cross-talk filters should therefore strongly favor minimizing FNs by default.

### Parameter tuning and validation

The UNCROSS2 algorithm has user-settable parameters *s, N*_*min*_, *S*_*min*_, *f*_*max*_, *C*_*min*_ and *t*_*min*_. Ideally, these would be trained and validated using several independent datasets with control samples. However, the only suitable training dataset I am aware of is Koz2013. I therefore used an alternative strategy, as follows. I selected default parameter values which seemed intuitively reasonable and produced similar results to my own manual analyses of OTU tables based on informed guesswork. To investigate whether results are robust against varying parameter values, I measured predicted cross-talk rates using all combinations of parameters shown in Table 2. Using the mock samples in Koz2013, I measured the number of true positives (TPs, i.e. non-zero counts which were correctly identified as solely due to cross-talk) true negatives, (TNs, i.e. counts which were correctly identified as valid non-zero values), FPs and FNs for each set of parameter values. Denoting the number of TPs as *NTP* etc., I calculated the sensitivity *Sens* = *NTP*/(*NTP* + *NFN*) and error rate *Err* = (*NFP* + *NFN*)/(*NTP* + *NTN* + *NFP* + *NFN*).

**Table 2.** Tested parameters for the UNCROSS2 algorithm. Defaults are highlighted. All of the 48 possible combinations obtained by selecting one value for each parameter were tested. The *de novo* rate estimate is not meaningfully sensitive to parameters *S*_*min*_ and *C*_*min*_, which were therefore excluded from parameter variation testing.

| *Parameter* | *Values* |
| --- | --- |
| $f_{max}$ | 0.01, **0.02**, 0.05, 0.1 |
| $s$ | 0.01, 0.05, **0.1**, 0.2 |
| $N_{min}$ | 100, 500, **1000** |

### OTU table coloring

To facilitate manual (i.e., visual) review, UNCROSS2 optionally generates an OTU table in HTML format where the cell for each count is colored according to the score given by eq. (3). A zero count is indicated by a blank white cell. If *t*≥0.5, the background color is dark orange, if *t*≥0.1 (the default threshold), the color is light orange, otherwise green. An example is shown in Fig. 2.

**Figure 2.**
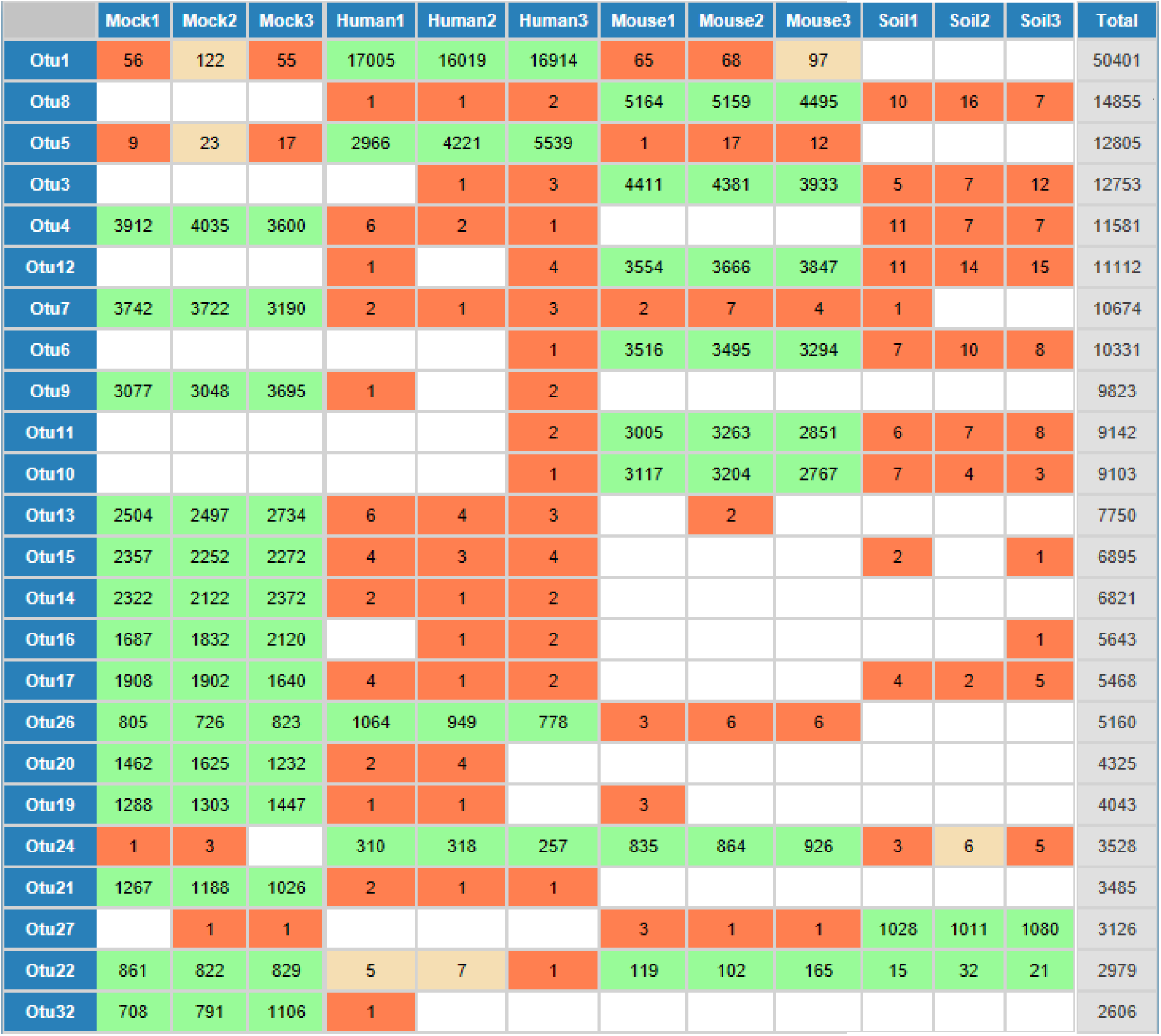
OTU table with counts colored by UNCROSS2 score. This is a partial table for Koz2013 run 134017 showing the most abundant OTUs. The cell for each count is colored according to the UNCROSS2 score given by eq. (3). A zero count is indicated by a blank white cell. If *t*≥0.5, the background color is dark orange, if *t*≥0.1 (the default threshold), the color is light orange, otherwise green.

## Results

### Accuracy on Koz2013 runs

Results for the eleven runs in Koz2013 are summarized in Table 3. Cross-talk rates measured on mock samples are in good agreement with rates estimated *de novo*, consistently reporting ∼1% mis-assigned reads. The largest disagreement is on run 130125, where the *de novo* estimate (0.013) is approximately twice as large as the measured mock rate (0.0068), noting that the true mean over all samples may differ from a subset such as mock samples. Sensitivity ranges from 77% (run 130306) to 89% (three different runs), while the error rate ranges from 1% (130125) to 20% (130306). Sensitivity in the range 77% to 89% would often be considered good performance for a bioinformatics algorithm, but here it corresponds to a false negative rate of 11% to 23% which could substantially inflate diversity estimates in practice.

**Table 3.** Results on the Koz2013 runs. These results were obtained using default parameters.

| *Run* | *Mock rate* | *De novo rate* | *TP* | *TN* | *FP* | *FN* | *Sens.* | *Err.* |
| --- | --- | --- | --- | --- | --- | --- | --- | --- |
| 121203 | 0.0063 | 0.0059 | 96 | 57 | 0 | 14 | 87% | 8% |
| 121205 | 0.0078 | 0.0065 | 191 | 57 | 0 | 28 | 87% | 10% |
| 121207 | 0.0075 | 0.0081 | 166 | 59 | 0 | 16 | 91% | 7% |
| 130125 | 0.0068 | 0.013 | 29 | 60 | 0 | 1 | 97% | 1% |
| 130211 | 0.0077 | 0.0075 | 125 | 60 | 0 | 22 | 85% | 11% |
| 130220 | 0.0083 | 0.0082 | 322 | 58 | 0 | 33 | 91% | 8% |
| 130306 | 0.015 | 0.0089 | 288 | 57 | 0 | 85 | 77% | 20% |
| 130401 | 0.015 | 0.011 | 615 | 57 | 0 | 78 | 89% | 10% |
| 130403 | 0.015 | 0.010 | 702 | 60 | 0 | 72 | 91% | 9% |
| 130417 | 0.014 | 0.0085 | 448 | 60 | 0 | 55 | 89% | 10% |
| 130422 | 0.013 | 0.0091 | 454 | 60 | 0 | 55 | 89% | 10% |

### Accuracy with varying threshold

Table 4 shows results obtained with varying *t*_*min*_ values on a typical Koz2013 run (130417). Both sensitivity and error rate improve with smaller values of *t*_*min*_, noting that there is some redundancy between the two measures because the error rate includes false negatives which are also reflected in the sensitivity. If parameters were tuned to this data, the smallest value of *t*_*min*_ would be selected. However, I believe this is an artifact of the mock community which has a much smaller number of OTUs than are typically found *in vivo*. In this artificial case, removing most small counts gives a more accurate OTU table, while in practice it would probably tend to give an unacceptably high false positive rate.

**Table 4.** Results on Koz2013 run 134017 obtained with varying *tmin* values. Accuracy improves with smaller *t*_*min*_, but this may be unrealistic in practice due to the low diversity of the mock samples (see main text for discussion).

| $t_{min}$ | $TP$ | $TN$ | $FP$ | $FN$ | $Sens$ | $Err$ |
| --- | --- | --- | --- | --- | --- | --- |
| 0.0001 | 485 | 60 | 0 | 18 | 96.42 | 3.2 |
| 0.001 | 479 | 60 | 0 | 24 | 95.23 | 4.26 |
| 0.01 | 468 | 60 | 0 | 35 | 93.04 | 6.22 |
| 0.1 | 448 | 60 | 0 | 55 | 89.07 | 9.77 |
| 0.5 | 385 | 60 | 0 | 118 | 76.54 | 20.96 |

### Robustness of de novo rate estimate against varying parameters

Results with varying parameters for all datasets are summarized in Table 5. On most datasets, the standard deviation of the *de novo* rate estimate is small compared to the mean, indicating it is robust against variations in the parameters. This also shows that parameters are not over-tuned to Koz2013, which is the only dataset for which cross-talk can be determined independently of the OTU table. The mean value over all parameter sets is similar to the default value on all datasets where a *de novo* estimate is reported.

**Table 5.** Results obtained with varying parameters. Here, *De novo rate* is the rate predicted with default parameters, *Avg. rate* is the mean *de novo* rate over all tested parameter values, and *Std. dev.* is the standard deviation of the rate. Cases where <10 candidate OTUs were found are undetermined (*undet*., *n* where *n* is the number of candidates). These results show that the estimated *de novo* rate is robust against varying parameters and ambiguous data, and that the parameters are not over-tuned to the Koz2013 dataset.

| *Dataset* | *Run* | *Samples* | *Mock rate* | *De novo rate* | *Avg. rate* | *Std. dev.* |
| --- | --- | --- | --- | --- | --- | --- |
| Koz2013 | 121203 | Mock+soil+gut | 0.0063 | 0.0059 | 0.0072 | 0.003 |
| Koz2013 | 121205 | Mock+soil+gut | 0.0078 | 0.0065 | 0.0088 | 0.005 |
| Koz2013 | 121207 | Mock+soil+gut | 0.0075 | 0.0081 | 0.0089 | 0.005 |
| Koz2013 | 130125 | Mock+soil+gut | 0.0068 | 0.013 | 0.013 | 0.0006 |
| Koz2013 | 130211 | Mock+soil+gut | 0.0077 | 0.0075 | 0.0083 | 0.004 |
| Koz2013 | 130220 | Mock+soil+gut | 0.0083 | 0.0082 | 0.0090 | 0.004 |
| Koz2013 | 130306 | Mock+soil+gut | 0.015 | 0.0089 | 0.0099 | 0.006 |
| Koz2013 | 130401 | Mock+soil+gut | 0.015 | 0.011 | 0.010 | 0.005 |
| Koz2013 | 130403 | Mock+soil+gut | 0.015 | 0.010 | 0.01 | 0.005 |
| Koz2013 | 130417 | Mock+soil+gut | 0.014 | 0.0085 | 0.0098 | 0.005 |
| Koz2013 | 130422 | Mock+soil+gut | 0.013 | 0.0091 | 0.01 | 0.006 |
| Car2016 | - | Soil | - | (undet., 0) | - | - |
| Dev2018 | - | AD plants | - | (undet., 1) | - | - |
| Gev2014 | - | Rectum+ileum+gut | - | 0.016 | 0.024 | 0.02 |
| Gil2016 | - | Gut | - | 0.015 | 0.021 | 0.02 |
| Mac2015 | - | Vagina | - | (undet., 5) | - | - |
| Now2015 | - | Gut | - | 0.016 | 0.023 | 0.02 |
| Par2017 | - | Phytoplankton bloom | - | (undet., 3) | - | - |
| Tho2016 | - | Gut | - | (undet., 8) | - | - |
| Vir2017 | - | Vagina | - | 0.012 | 0.014 | 0.01 |
| Yow2017 | V2-V3 | Prostate | - | 0.0048 | 0.0041 | 0.002 |
| Yow2017 | V4 | Prostate | - | 0.0065 | 0.0069 | 0.003 |

## Discussion

### De novo estimation of cross-talk rate

Several lines of evidence suggest that UNCROSS2 reports a good *de novo* estimate of the cross-talk rate: agreement with measurements on mock samples, robustness against varying parameters, and the observation that estimated cross-talk rates are comparable across several diverse datasets (Table 5), as would be expected on the assumptions that cross-talk errors occur with similar rates in different studies using similar sequencing protocols and are independent of the biological sequences in the reads.

### An attempt to filter cross-talk, with limited success

On the mock samples in the eleven Koz2013 runs, UNCROSS2 filtering has sensitivity 77% to 91% and an error rate of 1% to 20%. While these results suggest that the algorithm is reasonably effective in removing many of the spurious counts due to cross-talk, this may not be sufficient to decisively improve diversity estimates in practice.

### Cross-talk is probably ubiquitous in practice

Estimated cross-talk rates on the tested datasets range from 0.4% mis-assigned reads (Yow2017, V4) to1.6% (Gev2014). While it cannot be ruled out that *de novo* values are overestimated for reasons that are currently unknown, it is conservative to assume that a cross-talk rate of ∼1% is typical in practice.

### Better data is needed

Analysis of cross-talk is severely hampered by limitations in available data. Most studies do not include control samples, precluding reliable measurements of cross-talk (or other types of error). In the case of MiSeq, demultiplexing is performed by the Illumina platform software which generates separate FASTQ files for each sample. The underlying data used to perform demultiplexing, i.e. the index reads and their quality scores, are generally not provided to the user. Next-generation amplicon sequencing datasets in public archives such as the NCBI Short Read Archive and the European Nucleotide Archive rarely include usable information about index sequences, index reads or which samples were sequenced together in a single run.

### Possible improvements to the algorithm

UNCROSS2 analyses an OTU table without considering other information that is potentially informative such as a list of the index sequences assigned to each sample, index reads sequences and index read quality scores. This design choice is pragmatic: an OTU table is almost always available, or can be constructed from available data, while index reads are rarely available. It seems likely that accuracy could be improved by considering the index reads, but this would have limited value until such time as index reads are routinely provided by sequencing machine software and are routinely deposited in public archives to enable independent re-analysis of published datasets.

### Mitigating cross-talk by modified indexing schemes

Given that cross-talk with currently popular indexing schemes cannot be reliably filtered, the problem can be more effectively addressed by modifications to the PCR and sequencing protocol such as those proposed in (Kircher *et al.*, 2012; Esling *et al.*, 2015).

### Using UNCROSS2 in practice

When control samples are included, UNCROSS2 can provide an accurate measurement of cross-talk into the controls. When control samples are not available, its *de novo* estimates of the cross-talk rate appear to be reliable, though this is not definitively established by the results in this paper because *de novo* predictions could be verified only on one dataset (Koz2013). An accurate estimate of the overall mean cross-talk rate is useful for assessing the scope of the problem, but does not necessarily enable effective filtering because of fluctuations around expected values. With currently popular indexing protocols, cross-talk analysis should be performed on a separate OTU table for each sequencing run because cross-talk between samples in different runs cannot occur. Given that filtering may have a high error rate and that an optimal threshold is ill-defined and/or difficult to determine, I suggest that the most robust approach is to perform diversity analysis multiple times using tables filtered at different thresholds. Biological conclusions are supportable if they are repeatable across these tables, e.g. if significant *P*-values are obtained on all of them.

